# Peptide Gaussian accelerated molecular dynamics (Pep-GaMD): Enhanced sampling and free energy and kinetics calculations of peptide binding

**DOI:** 10.1101/2020.07.13.200774

**Authors:** Jinan Wang, Yinglong Miao

## Abstract

Peptides mediate up to 40% of known protein-protein interactions in higher eukaryotes and play an important role in cellular signaling. However, it is challenging to simulate both binding and unbinding of peptides and calculate peptide binding free energies through conventional molecular dynamics, due to long biological timescales and extremely high flexibility of the peptides. Based on the Gaussian accelerated molecular dynamics (GaMD) enhanced sampling technique, we have developed a new computational method “Pep-GaMD”, which selectively boosts essential potential energy of the peptide in order to effectively model its high flexibility. In addition, another boost potential is applied to the remaining potential energy of the entire system in a dual-boost algorithm. Pep-GaMD has been demonstrated on binding of three model peptides to the SH3 domains. Independent 1 μs dual-boost Pep-GaMD simulations have captured repetitive peptide dissociation and binding events, which enable us to calculate peptide binding thermodynamics and kinetics. The calculated binding free energies and kinetic rate constants agreed very well with available experimental data. Furthermore, the all-atom Pep-GaMD simulations have provided important insights into the mechanism of peptide binding to proteins that involves long-range electrostatic interactions and mainly conformational selection. In summary, Pep-GaMD provides a highly efficient, easy-to-use approach for unconstrained enhanced sampling and calculations of peptide binding free energies and kinetics.

**Significance Statement:** We have developed a new computational method “Pep-GaMD” for enhanced sampling of peptide-protein interactions based on the Gaussian accelerated molecular dynamics (GaMD) technique. Pep-GaMD works by selectively boosting the essential potential energy of the peptide to effectively model its high flexibility. In addition, another boost potential can be applied to the remaining potential energy of the entire system in a dual-boost algorithm. Pep-GaMD has been demonstrated on binding of three model peptides to the SH3 domains. Dual-boost Pep-GaMD has captured repetitive peptide dissociation and binding events within significantly shorter simulation time (microsecond) than conventional molecular dynamics. Compared with previous enhanced sampling methods, Pep-GaMD is easier to use and more efficient for unconstrained enhanced sampling of peptide binding and unbinding, which provides a novel physics-based approach to calculating peptide binding free energies and kinetics.

## Introduction

Peptides mediate up to 40% of known protein-protein interactions in higher eukaryotes^1^. They play a key role in cellular signaling, protein trafficking, immune response and oncology^1, 2^. Protein-peptide interactions have emerged as attractive drug targets for both small molecules and designed inhibitory peptides^3–5^. Increasing number of peptide-based drugs is being licensed to market in recent years^6–9^. Therefore, understanding molecular mechanism of protein-peptide recognition has important applications in the fields of biology, medicine, and pharmaceutical sciences.

It is important to characterize protein-peptide binding conformations for rational structurebased design of peptide drugs. Experimental techniques including X-ray crystallography, nuclear magnetic resonance (NMR) and cryo-electron microscopy (cryo-EM) have been utilized to determine the structures of protein-peptide complexes^2^. Recent years have seen dramatic increase in the number of experimental protein-peptide structures. However, only static snapshots of the protein-peptide interactions could usually be captured in these structures. The thermodynamics and dynamic mechanism of peptide recognition by proteins remains poorly understood^10^.

Computational methods including peptide docking have been developed to model peptide-protein structures^11, 12^. In this regard, modeling of peptide binding to proteins has been shown to be distinct from that of extensively studied protein-ligand binding and protein-protein interactions. Notably, small-molecule ligands are able to bind deeply buried sites in proteins, but peptides normally bind to protein surface, especially in the large pockets. On the other hand, peptide-protein interactions are typically weaker than protein-protein interactions, because of the small interface between peptides and their target proteins. Moreover, protein partners usually have well defined 3D structures before forming protein-protein complexes, despite possible conformational changes during association. In contrast, most peptides do not have stable structures before forming complexes with proteins. It is difficult to incorporate extremely high flexibility of peptides and their large conformational changes (folding and unfolding) into computational modeling, especially for peptide docking^11^.

Molecular dynamics (MD) is a powerful technique for all-atom simulations of biomolecules^13^. MD has been used to refine binding poses of peptides in proteins obtained from docking^14–18^. In addition, MD simulations are able to account for the flexibility of peptides and have been applied to explore their mechanisms of binding to proteins^8, 19–26^. Notably, MD simulations have successfully revealed the pathway of fast association of a proline-rich motif (PRM) peptide to the SH3 domain^19^. Supervised MD simulations have captured binding pathways of a natural peptide BAD to the Bc1-X_L_ protein and the p53 peptide and SAH-p53-8 stapled peptidomimetic to the MDM2 protein^24^. Weighted ensemble of a total amount of ~120 μs MD simulations has been obtained to investigate binding of the p53 peptide to the MDM2 protein in implicit solvent^23^. The simulation predicted binding rate constant agrees very well with the experiments. Very recently, MD simulation performed for 200 μs at elevated temperature (400 K) using the Anton specialized supercomputer has captured 70 binding and unbinding events between an intrinsically disordered protein (IDP) fragment of the measles virus nucleoprotein and the X domain of the measles virus phosphoprotein complex, which enables detailed understanding of the peptide “folding-upon-binding” mechanism^26^. Despite these remarkable advances, these studies based on conventional MD (cMD) often suffer from high computational cost for simulations of peptide-protein interactions. It remains challenging to sufficiently sample peptide-protein interactions through cMD simulations, especially for slow dissociation of various peptides that could take place over milliseconds and longer timescales^27–31^.

Furthermore, enhanced sampling MD methods, including steered MD^32^, Modeling by Employing Limited Data (MELD) using temperature and Hamiltonian replica exchange MD^33, 34^, temperature-accelerated MD^35^, multi-ensemble Markov state models^36^ and metadynamics^37^, have been applied to improve sampling of peptide-protein interactions. Steered MD have allowed simulations of antigenic peptide unbinding from the T-cell receptor based on a dissociation reaction coordinate^32^. MELD-accelerated MD simulations have been performed to predict binding poses and relative binding free energies of the p53-derived peptides and stapled α-helical peptides to the MDM2 and MDMX proteins^33, 34^. Multi-ensemble Markov models combining hundreds-of-microsecond cMD and Hamiltonian replica exchange simulations have been implemented to characterize both dissociation and binding of the PMI peptide inhibitor to the MDM2 oncoprotein fragment^36^. While cMD was able to simulate fast events such as peptide binding, enhanced sampling simulations could capture rare events such as peptide unbinding beyond the seconds timescale. The simulation derived dissociation and association rates were found in good agreement with the experimental measurements^36^. Recently, bias-exchange metadynamics simulations with 8 defined collective variables (CVs) performed for a total of 27 μs have been used to calculate binding free energy of the p53 peptide to the MDM2 protein. Infrequent metadynamics simulations with 3 CVs have also been successfully applied to predict peptide binding and dissociation rates for the same system^37^. Therefore, enhanced MD simulations have greatly expanded our capabilities in studies of peptide-protein interactions. Nevertheless, enhanced sampling of peptide binding to proteins are still underexplored compared with more extensive studies of protein-ligand binding and protein-protein interactions^38–44^. The current enhanced sampling approaches are still computationally expensive for characterizing peptide binding thermodynamics and kinetics, requiring tens to hundreds of microsecond simulations.

Here, we will develop a new, easy-to-use enhanced sampling approach based on the robust Gaussian accelerated molecular dynamics (GaMD) technique for highly efficient and accurate simulations of peptide-protein interactions. GaMD works by adding a harmonic boost potential to smooth biomolecular potential energy^45^. It greatly reduces system energy barriers and accelerates biomolecular simulations by orders of magnitude^45^–^48^. GaMD provides unconstrained enhanced sampling without the requirement of predefined reaction coordinates or CVs. Compared with enhanced sampling methods that rely on careful selection of CVs, GaMD could be much easier to use for studying complex biological processes such as peptide binding to proteins^44^. Moreover, because the boost potential follows a Gaussian distribution, biomolecular free energy profiles can be properly recovered through cumulant expansion to the second order.^45^

Based on GaMD, we have recently developed a new ligand GaMD or “LiGaMD” method, which allows us to characterize both thermodynamics and kinetics of ligand binding quantitatively^49^. In LiGaMD, we selectively boost the ligand non-bonded interaction potential energy to enable ligand dissociation. Another boost potential is applied to the remaining potential energy of the entire system in a dual-boost algorithm to facilitate ligand rebinding. LiGaMD has been demonstrated on host-guest and protein-ligand binding model systems. Repetitive guest binding and unbinding in the β-cyclodextrin host were observed in hundreds-of-nanosecond LiGaMD simulations. The calculated guest binding free energies agreed excellently with experimental data with < 1.0 kcal/mol errors. Compared with converged microsecond-timescale cMD simulations, the sampling errors of LiGaMD_Dual simulations were also < 1.0 kcal/mol. Accelerations of ligand kinetic rate constants in LiGaMD simulations were properly estimated using Kramers’ rate theory. Furthermore, LiGaMD allowed us to capture repetitive dissociation and binding of the benzamidine inhibitor in trypsin within 1 μs simulations. The calculated ligand binding free energy and kinetic rate constants compared very well with the experimental data^49^.

Building upon GaMD and LiGaMD, we present a new computational method called peptide GaMD (“Pep-GaMD”), which enables us to simulate repetitive peptide binding and unbinding within microsecond simulations. In Pep-GaMD, the essential potential energy of the peptide is selectively boosted to effectively model its high flexibility. Selective acceleration has been found useful in previous enhanced sampling techniques, including selective aMD^50^, selectively scaled MD^51^, essential energy space random walk^52, 53^ and LiGaMD^49^. In addition, another boost potential is applied on the protein and solvent in a dual-boost Pep-GaMD algorithm. To demonstrate the new Pep-GaMD method, the SH3 domains with important biological functions^54, 55^ is selected as our model system. Pep-GaMD has been testing on the binding of three PRM peptides, including “PAMPAR” (PDB: 1SSH), “PPPALPPKK” (PDB: 1CKA) and “PPPVPPRR” (PDB: 1CKB). Repetitive binding and unbinding of the three peptides to the SH3 domains have been observed in microsecond Pep-GaMD simulations. The peptide binding free energies and kinetic rate constants calculated from Pep-GaMD simulations agree very well with available experimental data. Furthermore, the Pep-GaMD simulations have provided important insights into the binding mechanism of peptide to their target proteins at an atomistic level.

## Methods

### Peptide Gaussian accelerated molecular dynamics (Pep-GaMD)

Gaussian accelerated molecular dynamics (GaMD) is an enhanced sampling technique that works by adding a harmonic boost potential to smooth biomolecular potential energy surface and reduce the system energy barriers^45^. Details of the GaMD method have been described in previous studies^45–47^. A brief summary is provided in **Supplementary Material**. Here, we developed a new Pep-GaMD method for more efficient sampling of peptide binding to proteins.

We consider a system of peptide *L* binding to a protein *P* in a biological environment *E*. The system comprises of *N* atoms with their coordinates 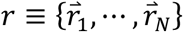 and momenta 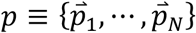. The system Hamiltonian can be expressed as:

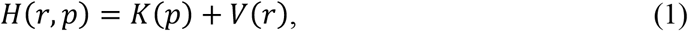

where *K(p)* and *V(r)* are the system kinetic and total potential energies, respectively. Next, we decompose the potential energy into the following terms:

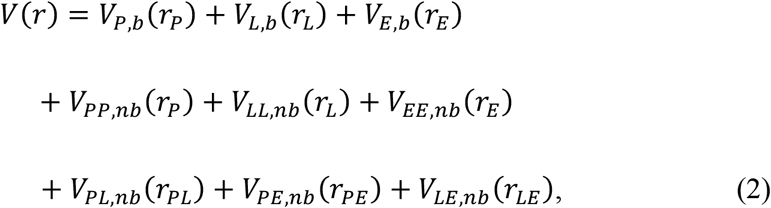

where *V_P,b_*, *V_L,b_* and *V_E,b_* are the bonded potential energies in protein *P*, peptide *L* and environment *E*, respectively. *V_PP,nb_*, *V_LL,nb_* and *V_EE,nb_* are the self non-bonded potential energies in protein *P*, peptide *L* and environment *E*, respectively. *V_PL,nb_*, *V_PE,nb_* and *V_LE,nb_* are the non-bonded interaction energies between *P-L, P-E* and *L-E*, respectively. According to classical molecular mechanics force fields^56, 57^, the non-bonded potential energies are usually calculated as:

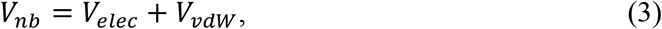

where *V_elec_* and *V_vdw_* are the system electrostatic and van der Waals potential energies. Presumably, peptide binding mainly involves in both the bonded and non-bonded interaction energies of the peptide since peptides often undergo large conformational changes during binding to the target proteins. Thus, the essential peptide potential energy is *V_L_(r) = V_LL,b_ (r_L_)* + *V_LL,nb_ (r_L_)* + *VpL,nb (r_PL_)* + *V_LE,nb_ (r_LE_)*. In Pep-GaMD, we add boost potential selectively to the essential peptide potential energy according to the GaMD algorithm:

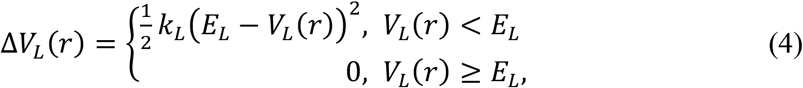

where *E_L_* is the threshold energy for applying boost potential and *K_L_* is the harmonic constant. The Pep-GaMD simulation parameters are derived similarly as in the previous GaMD algorithm (see details in **Supplementary Material**). When *E* is set to the lower bound as the system maximum potential energy (*E*=*V_max_*), the effective harmonic force constant *k*_0_ can be calculated as:

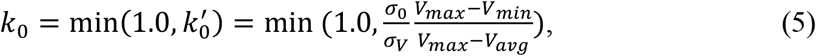

where *V_max_, V_min_, V_avg_* and *σ_v_* are the maximum, minimum, average and standard deviation of the boosted system potential energy, and *σ*_0_ is the user-specified upper limit of the standard deviation of *ΔV* (e.g., 10*K_B_T*) for proper reweighting. The harmonic constant is calculated as 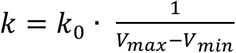 with 0 < *k*_0_ ≤ 1. Alternatively, when the threshold energy *E* is set to its upper bound 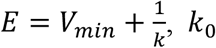 is set to:

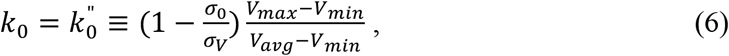

if 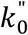 is found to be between *0* and *1*. Otherwise, *k*_0_ is calculated using Eqn. (5).

In addition to selectively boosting the peptide, another boost potential is applied on the protein and solvent to enhance conformational sampling of the protein and facilitate peptide binding. The second boost potential is calculated using the total system potential energy other than the essential peptide potential energy as:

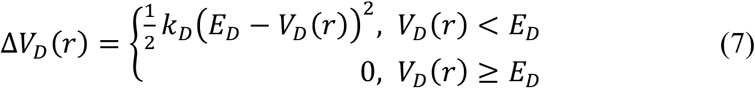

Where V_D_ is the total system potential energy other than the essential peptide potential energy, E_D_ is the corresponding threshold energy for applying the second boost potential and k_D_ is the harmonic constant. This leads to dual-boost Pep-GaMD with the total boost potential Δ*V(r)* = *ΔV_L_ (r)* + *ΔV_D_ (r)*. Pep-GaMD is currently implemented in the GPU version of AMBER 20^58^, but should be transferable to other MD programs as well (see **Supplementary Material**).

The Pep-GaMD approach is in contrast to previous standard GaMD, in which the boost potential was applied to the system dihedrals and/or total potential energy^45–47^. It is based on earlier studies showing that the essential energy space random walk provided very efficient enhanced sampling^52, 53, 59^ and convergence of aMD simulations was improved by selectively applying boost potential to only interesting regions of biomolecules^50^. Pep-GaMD also uses distinct boost potential formulas compared with LiGaMD, in which only the non-bonded potential energy of a bound ligand was selectively boosted to accelerate its dissociation^49^. In comparison, the total potential energy of both bonded and non-bonded interactions is boosted in Pep-GaMD for a peptide that is significantly more flexible than a small-molecule ligand.

### Peptide binding free energy calculations from 3D potential of mean force

We calculate peptide binding free energy from 3D potential of mean force (PMF) of peptide displacements from the target protein as the following^60, 61^:

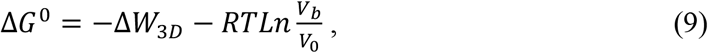

where *V*_0_ is the standard volume, *V_b_* = ∫ *e^-βW(r)^ dr* is the average sampled bound volume of the peptide with *β* = 1/*k_B_T, k_B_* is the Boltzmann constant, *T* is the temperature, and *ΔW_3D_* is the depth of the 3D PMF. *ΔW_3D_* can be calculated by integrating Boltzmann distribution of the 3D PMF *W(r)* over all system coordinates except the x, y, z of the peptide:

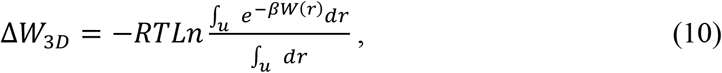

where *V_u_ = ∫_U_ dr* is the sampled unbound volume of the peptide. The exact definitions of the bound and unbound volumes *V_b_* and *V_u_* are not important as the exponential average cut off contributions far away from the PMF minima^61^. A python script “PyReweighting-3D.py” in the *PyReweighting* tool kit http://miao.compbio.ku.edu/PyReweighting/)^49, 62^ was applied for reweighting Pep-GaMD simulations (see Methods in **Supplementary Material**) to calculate the 3D reweighted PMF and associated peptide binding free energies.

### Peptide binding kinetics obtained from reweighting of Pep-GaMD Simulations

Reweighting of peptide binding kinetics from Pep-GaMD simulations followed a similar protocol using Kramers’ rate theory that has been recently implemented in kinetics reweighting of the GaMD^48^ and LiGaMD^49^ simulations. Provided sufficient sampling of repetitive peptide dissociation and binding in the simulations, we record the time periods and calculate their averages for the peptide found in the bound (*τ_B_*) and unbound (*τ_u_*) states from the simulation trajectories. The *τ_B_* corresponds to residence time in drug design^63^. Then the peptide dissociation and binding rate constants (*k*_off_ and *k*_on_) were calculated as:

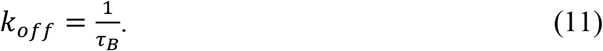

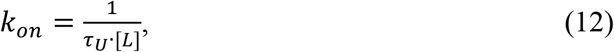

where [L] is the peptide concentration in the simulation system.

According to Kramers’ rate theory, the rate of a chemical reaction in the large viscosity limit is calculated as^48^:

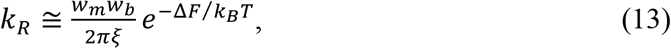

where *w_m_* and *w_b_* are frequencies of the approximated harmonic oscillators (also referred to as curvatures of free energy surface^64, 65^) near the energy minimum and barrier, respectively, ξ is the frictional rate constant and *ΔF* is the free energy barrier of transition. The friction constant *ξ* is related to the diffusion coefficient *D* with *ξ* = *k_B_T/D*. The apparent diffusion coefficient *D* can be obtained by dividing the kinetic rate calculated directly using the transition time series collected directly from simulations by that using the probability density solution of the Smoluchowski equation^66^. In order to reweight peptide kinetics from the Pep-GaMD simulations using the Kramers’ rate theory, the free energy barriers of peptide binding and dissociation are calculated from the original (reweighted, ***ΔF***) and modified (no reweighting, ***ΔF***\*) PMF profiles, similarly for curvatures of the reweighed (*w*) and modified (*w*\*, no reweighting) PMF profiles near the peptide bound (“B”) and unbound (“U”) low-energy wells and the energy barrier (“Br”), and the ratio of apparent diffusion coefficients from simulations without reweighting (modified, *D*\*) and with reweighting (*D*). The resulting numbers are then plugged into Eq. (13) to estimate accelerations of the peptide binding and dissociation rates during Pep-GaMD simulations^48^, which allows us to recover the original kinetic rate constants.

### Protein-peptide binding simulations

Pep-GaMD simulations using the dual-boost scheme were performed on binding of three model peptides to the SH3 domains. X-ray crystal structures of the SH3 domains bound by “PAMPAR” (PDB: 1SSH), “PPPALPPKK” (PDB: 1CKA)^67^ and “PPPVPPRR” (PDB: 1CKB)^67^ were used.

The N-and C-termini of the peptides were capped with the acetyl (ACE) and primary amide (NHE) neutral groups, respectively. The missing hydrogen atoms were added using the *tleap* module in AMBER^68^. The AMBER ff14SB force field^69^ were used for the peptides and protein. Each system was neutralized by adding counter ions and immersed in a cubic TIP3P water box^70^, which was extended 13 Å from the protein-peptide complex surface.

Each simulation system was first energy minimized with 1.0 kcal/mol/Å^2^ constraints on the heavy atoms of the protein and peptide, including the steepest descent minimization for 50,000 steps and conjugate gradient minimization for 50,000 steps. The system was then heated from 0 K to 300 K for 200 ps. It was further equilibrated using the NVT ensemble at 300K for 200 ps and the NPT ensemble at 300 K and 1 bar for 1 ns with 1 kcal/mol/Å^2^ constraints on the heavy atoms of the protein and peptide, followed by 2 ns short cMD without any constraint. The Pep-GaMD simulations proceeded with 14 ns short cMD to collect the potential statistics, 46 ns Pep-GaMD equilibration after adding the boost potential and then three independent 1000 ns production runs.

It provided more powerful sampling to set the threshold energy for applying the boost potential to the upper bound (i.e., *E* = *V*_min_+1/*k*) in our previous study ligand dissociation and binding using LiGaMD^49^. Therefore, the threshold energy for applying the peptide boost potential was also set to the upper bound in the Pep-GaMD simulations. For the second boost potential applied on the system total potential energy other than the essential peptide potential energy, sufficient acceleration was obtained by setting the threshold energy to the lower bound. In order to observe peptide dissociation during Pep-GaMD equilibration while keeping the boost potential as low as possible for accurate energetic reweighting, the (σ_0P_, σ_0D_) parameters were finally set to (2.2 kcal/mol, 6.0 kcal/mol), (4.0 kcal/mol, 6.0 kcal/mol) and (4.0 kcal/mol, 6.0 kcal/mol) for the Pep-GaMD simulations of the 1SSH, 1CKA and 1CKB systems, respectively. Pep-GaMD production simulation frames were saved every 0.2 ps for analysis.

The VMD^71^ and CPPTRAJ^72^ tools were used for simulation analysis. The 1D, 2D and 3D PMF profiles, as well as the peptide binding free energy, were calculated through energetic reweighting of the Pep-GaMD simulations. Root-mean square deviations (RMSDs) of the peptides relative to X-ray structures with the protein aligned, the peptide radius of gyration (*Rg*), distances between two protein residues that are located in the peptide-binding sites (denoted *d*_D-W_ for Asp19-Trp40 in the 1SSH structure and Asp150-Trp169 in the 1CKA and 1CKB structures) and a salt bridge distance between the protein and peptides (denoted d_D-R/K_ for Asp19 (SH3)-Arg10 (peptide) in the 1SSH structure, Asp150 (SH3)-Lys8 (peptide) in the 1CKA structure and Asp150 (SH3)-Arg7 (peptide) in the 1CKB structure) were chosen as reaction coordinates for calculating the PMF profiles. The RMSD of peptides with protein aligned was chosen as reaction coordinate for calculating the 1D PMF profile. The bin size was set to 1.0 Å. 2D PMF profiles of d_D-w_/d_D-R/K_/*R_g_* and peptide backbone RMSD were calculated to analyze conformational changes of the protein and important interactions upon peptide binding. The bin size was set to 1.0 Å for these reaction coordinates. The cutoff for the number of simulation frames in one bin was set to 500 for reweighting 1D and 2D PMF profiles. The 3D PMF profiles of peptide center-of-mass displacements from the SH3 domains in the X, Y and Z directions were further calculated from the Pep-GaMD simulations. The bin sizes were set to 1.0 Å in the X, Y and Z directions. The cutoff of simulation frames in one bin for 3D PMF reweighting was set to the minimum number below which the calculated global minimum of 3D PMF will be shifted. The peptide binding free energies (*ΔG*) were calculated using the reweighted 3D PMF profiles.

## Results and Discussions

### Microsecond Pep-GaMD simulations captured repetitive dissociation and binding of peptides to the SH3 domains

All-atom Pep-GaMD simulations were performed on X-ray crystal structures of the SH3 domains bound by three peptides, including “PAMPAR” (PDB: 1SSH), “PPPALPPKK” (PDB: 1CKA)^67^ and “PPPVPPRR” (PDB: 1CKB)^67^ (**Figs. 1A–1C**). Three independent 1000 ns Pep-GaMD production trajectories were obtained on each of the three peptide systems (**Table 1**). The Pep-GaMD simulations of the 1SSH system recorded an average boost potential of 12.83-13.01 kcal/mol with 3.75-3.80 kcal/mol standard deviation. In comparison, the average boost potential was 25.59-26.56 kcal/mol with 4.91-4.95 kcal/mol standard deviation in three simulations of the 1CKA system. The boost potential applied in simulations of the 1CKB system was similar to that of the 1CKA system, with average of 27.85-28.30 kcal/mol and 4.96-5.14 kcal/mol standard deviation (**Table 1**).

**Fig. 1.**
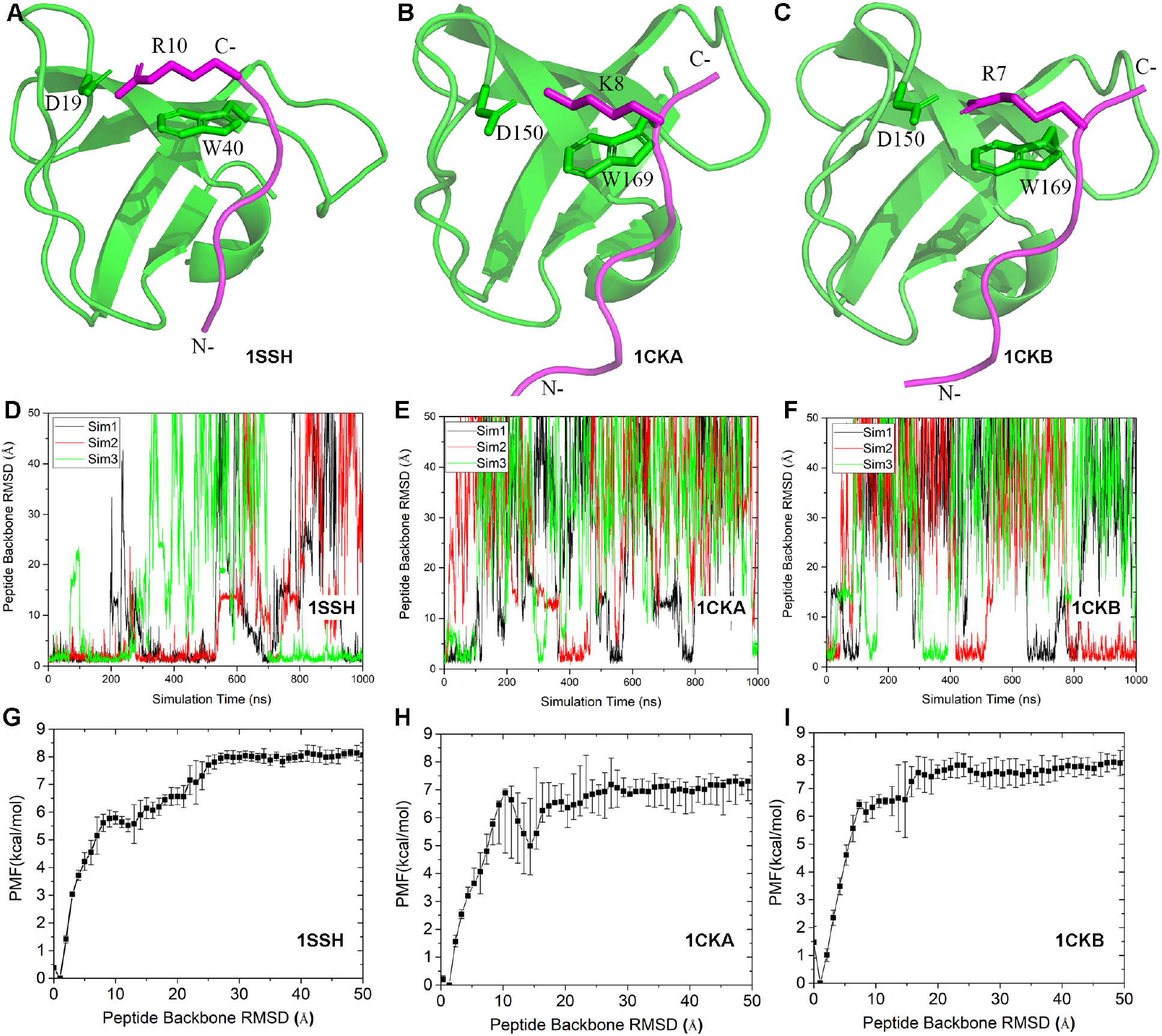
Pep-GaMD simulations have captured repetitive dissociation and binding of three model peptides to the SH3 domains: (A-C) X-ray structures of the SH3 domains bound by peptides (A) “PAMPAR” (PDB: 1SSH), (B) “PPPALPPKK” (PDB: 1CKA) and (C) “PPPVPPRR” (PDB: 1CKB). The SH3 domains and peptides are shown in green and magenta cartoon, respectively. Key protein residues Asp19 and Trp40 in the 1SSH structure and Asp150 and Trp169 in the 1CKA and 1CKB structures, and peptide residues Arg10 in the 1SSH structure, Lys8 in the 1CKA structure and Arg7 in the 1CKB structure are highlighted in sticks. The “N” and “C” labels denote the N-terminus and C-terminus of the peptides. (D-F) time courses of peptide backbone RMSDs relative to X-ray structures with the protein aligned calculated from three independent 1 μs Pep-GaMD simulations of the (D) 1SSH, (E) 1CKA and (F) 1CKB structures. (G-I) The corresponding PMF profiles of the peptide backbone RMSDs averaged over three Pep-GaMD simulations of the (G) 1SSH, (H) 1CKA and (I) 1CKB structures. Error bars are standard deviations of the free energy values calculated from three Pep-GaMD simulations.

**Table 1.** Summary of Pep-GaMD simulations performed on binding of three model peptides to the SH3 domains. *ΔV* is the total boost potential. *N_D_* and *N_B_* are the number of observed peptide dissociation and binding events, respectively. *ΔG_sim_* and *ΔG_exp_* are the peptide binding free energies obtained from Pep-GaMD simulations and experiments, respectively.

| PDB | ID | Length (ns) | $\Delta V$ (kcal/mol) | $N_D$ | $N_B$ | $\Delta G_{sim}$ (kcal/mol) | $\Delta G_{exp}^{64}$ (kcal/mol) |
| --- | --- | --- | --- | --- | --- | --- | --- |
| 1SSH | Sim1 | 1000 | 13.01±3.79 | 3 | 4 | -7.41±0.12 | - |
| | Sim2 | 1000 | 12.88±3.75 | 3 | 3 | | |
| | Sim3 | 1000 | 12.83±3.80 | 3 | 3 | | |
| 1CKA | Sim1 | 1000 | 25.65±4.91 | 4 | 5 | -7.72±0.54 | -7.84 |
| | Sim2 | 1000 | 26.56±4.95 | 3 | 3 | | |
| | Sim3 | 1000 | 25.59±4.89 | 2 | 3 | | |
| 1CKB | Sim1 | 1000 | 27.98±5.02 | 3 | 4 | -6.84±0.14 | -7.24 |
| | Sim2 | 1000 | 28.30±5.14 | 2 | 3 | | |
| | Sim3 | 1000 | 27.85±4.96 | 3 | 3 | | |

The peptide backbone RMSDs with the SH3 protein domain aligned were calculated as a function of simulation time (**Figs. 1D–1F**) to record the number of peptide dissociation and binding events (*N_D_* and *N_B_*) in each of the 1 μs Pep-GaMD simulations. With close examination of the peptide binding trajectories, cutoffs of the peptide backbone RMSD for the unbound and bound states were set to >25 Å and <5.0 Å, respectively. Because of the peptide fluctuations, we recorded only the corresponding events lasted for more than 1.0 ns. Repetitive dissociation and binding of the three peptides were successfully captured in each of the 1 μs Pep-GaMD simulations (**Figs. 1D–1F, S1**). In each simulation of the 1SSH system, about 3-4 binding and 3 dissociation events were observed for the peptide. In simulations of the 1CKA system, about 2-5 dissociation and 3-5 binding events were observed. A similar number of peptide dissociation (2-3) and binding (3-4) events were observed in simulations of the 1CKB system (**Table 1**).

Next, we computed 1D PMF free energy profiles from the Pep-GaMD simulations to quantitatively characterize peptide binding to the SH3 domains. The RMSD of each peptide relative to the native X-ray structure with the protein domain aligned was chosen as the reaction coordinate. The PMF profiles were calculated for three individual 1 μs Pep-GaMD simulations and averaged for each system (**Figs. 1G–1I**). The global free energy minima were all identified in the “Bound” state with ~2 Å peptide backbone RMSD. Error bars (standard deviations) of free energy values in the PMF profiles were mostly <1.0 kcal/mol, suggesting reasonable convergence of the Pep-GaMD simulations. Notably, differences were identified among the three peptide binding systems in the magnitudes of PMF values near the “Intermediate”, “Unbound” and energy barrier regions. The “Intermediate” low-energy state was identified for the 1SSH and 1CKA systems with the peptide backbone RMSD in the range of ~9-25 Å, although fluctuations were observed in the calculated PMF values (**Figs. 1G** and **1H**). In comparison, no clear low-energy well was identified in the “Intermediate” state for the 1CKB system, which showed rather a plateau in the region of 7.0 Å-25.0 Å peptide backbone RMSD (**Fig. 1I**). This plateau suggested no significant energy barrier and thus faster kinetic rate for peptide binding in the 1CKB system than in the other two systems (see details below). PMF values of the “Unbound” state relative to the “Bound” state were approximately 8.0 kcal/mol, 7.1 kcal/mol and 7.5 kcal/mol for three peptides in the 1SSH, 1CKA and 1CKB systems, respectively.

### Peptide binding free energies calculated from Pep-GaMD simulations agreed well with available experimental data

We computed binding free energies of three peptides to the SH3 domains based on their 3D PMF profiles (see details in **Methods**). The 3D PMF was calculated from each individual 1 μs Pep-GaMD simulation of peptide binding to the SH3 domain in terms of displacements of the Cα atom in the central peptide proline residue (Pro8 in the 1SSH structure, Pro6 in the 1CKA structure and Pro5 in the 1CKB structure) from the Cα atom in a protein residue (Asn55 in the 1SSH structure, Pro185 in the 1CKA and 1CKB structures) in the X, Y and Z directions. The 3D PMF was energetically reweighted through cumulant expansion to the second order (**Supplementary Material**). We then calculated the peptide binding free energies using the 3D reweighted PMF profiles. For the 1CKA system, the average of calculated peptide binding free energy values was −7.72 kcal/mol and the standard deviation was 0.54 kcal/mol, which was highly consistent with the experimental value of −7.84 kcal/mol^67^ (**Table 1**). For the 1CKB system, the resulting binding free energy was −6.84±0.14 kcal/mol, being closely similar to the experimental value of −7.24 kcal/mol^67^ (**Table 1**). In this context, not only the relative ranks of the peptide binding free energies were consistent with experimental data, the absolute binding free energy values calculated from Pep-GaMD simulations were also in excellent agreements with those from the experiments. The differences between the simulation predicted and experimentally determined binding free energies were less than 1.0 kcal/mol. An error of only 1.0 kcal/mol was widely used as a high level of prediction accuracy in small-molecule ligand binding free energy calculations^73, 74^. Provided the fact that peptides usually have larger sizes and significantly higher flexibility than the smallmolecule ligands, it would be valuable to achieve similar accuracy (1.0 kcal/mol error) in Pep-GaMD simulation prediction of the peptide binding free energy. For the 1SSH system, the calculated peptide binding free energy was −7.41±0.12 kcal/mol. However, experimental peptide binding free energy of this system was not available in the literature for comparison. According to the simulation prediction, the peptide binding free energy in the 1SSH system was between those of the 1CKA and 1CKB systems.

In summary, peptide binding free energies in the SH3 domains calculated from microsecond Pep-GaMD simulations agreed well with available experimental data. Compared with the experimental values, errors in the peptide binding free energies predicted from the Pep-GaMD simulations were smaller than 1.0 kcal/mol. Therefore, both efficient enhanced sampling and accurate free energy calculations of peptide binding were achieved through the Pep-GaMD simulations.

### Kinetics of peptide binding to SH3 domains

With accurate prediction of the peptide binding free energy, we analyzed the Pep-GaMD simulations further to determine the kinetic rate constants of peptide binding to SH3 domains. We recorded the time periods for the peptide found in the bound (τ_B_) and unbound (τ_U_) states throughout the Pep-GaMD simulations (**Table S1**). With only one peptide molecule in the simulations, the peptide concentrations were 0.0033 M, 0.0034 M and 0.0034 M in the 1SSH, 1CKA and 1CKB systems, respectively. Without reweighting of the Pep-GaMD simulations, the peptide binding (*k_on_*\*) and dissociation (*k_off_*\*) rate constants of 1SSH were calculated as 4.90 ± 0.73 × 10^10^ M^−1^·s^−1^ and 7.09 ± 1.27 × 10^6^ s^−1^, respectively. For 1CKA, the peptide binding (*k_on_*\*) and dissociation (*k_off_*\*) rate constants were calculated as 1.24 ± 0.27 × 10^9^ M^−1^·s^−1^ and 2.09 ± 0.44 × 10^7^ s^−1^, respectively. The peptide binding (*k_on_*\*) and dissociation (*k_off_*\*) rate constants in 1CKB were calculated as 1.81 ± 0.86 × 10^9^ M^−1^·s^−1^ and 1.28 ± 0.30 × 10^7^ s^−1^, respectively (**Table 2**).

**Table 2.** Comparison of kinetic rates obtained from experimental data and Pep-GaMD simulations for the three model peptides binding to the SH3 domains. *k_on_* and *k_off_* are the kinetic dissociation and binding rate constants, respectively, from experimental data or Pep-GaMD simulations with reweighting using Kramers’ rate theory. *k_on_*\* and *k_off_*\* are the accelerated kinetic dissociation and binding rate constants calculated directly from Pep-GaMD simulations without reweighting.

| | Method | $k_{on}$<br>($M^{-1} \cdot s^{-1}$) | $k_{off}$<br>($s^{-1}$) | $k_{on}^*$<br>($M^{-1} \cdot s^{-1}$) | $k_{off}^*$<br>($s^{-1}$) |
| --- | --- | --- | --- | --- | --- |
| 1SSH | Experiment | - | - | - | - |
| | Pep-GaMD | $1.92 \pm 1.17 \times 10^{10}$ | $749.19 \pm 1.57$ | $4.90 \pm 0.73 \times 10^{10}$ | $7.10 \pm 1.27 \times 10^6$ |
| 1CKA | Experiment | - | - | - | - |
| | Pep-GaMD | $1.80 \pm 1.05 \times 10^9$ | $511.65 \pm 314.04$ | $1.24 \pm 0.27 \times 10^9$ | $2.09 \pm 0.44 \times 10^7$ |
| 1CKB | Experiment<sup>76</sup> | $1.5 \times 10^9$ | $8.9 \times 10^3$ | - | - |
| | Pep-GaMD | $4.06 \pm 2.26 \times 10^{10}$ | $1.45 \pm 1.17 \times 10^3$ | $1.81 \pm 0.86 \times 10^9$ | $1.28 \pm 0.30 \times 10^7$ |

For 1CKA, the dissociation free energy barrier (Δ*F_off_*) decreased by ~81% from 7.42 ± 0.64 kcal/mol in the reweighted PMF profile to 1.35 ± 0.084 kcal/mol in the modified PMF profile (**Table S2**). The free energy barrier for peptide binding (Δ*F_on_*) decreased from 2.02 ± 1.12 kcal/mol in the reweighted profile to 1.17 ± 0.04 kcal/mol in the modified PMF profile (**Table S2**). According to the Kramers’ rate theory, the peptide binding rate was actually decreased by a factor of 0.69, while the peptide dissociation was accelerated by 4.08 × 10^4^ times. Therefore, the reweighted *k_on_* and *k_off_* were calculated as 1.80 ± 1.05×10^9^ M^-1^·s^-1^ and 511.65 ± 314.04 s^-1^, respectively. For 1CKB, the dissociation free energy barrier (Δ*F_off_*) decreased by ~79% from 7.84

Following a similar protocol as described in analysis of LiGaMD simulations^49^, we reweighted the peptide-SH3 domain binding simulations to calculate acceleration factors of the peptide binding and dissociation processes (**Table S2**) and recover the original kinetic rate constants using the Kramers’ rate theory. For 1SSH, the dissociation free energy barrier (Δ*F_off_*) decreased by ~70% from 8.16 ± 0.16 kcal/mol in the reweighted PMF profile to 2.40 ± 0.32 kcal/mol in the modified PMF profile (**Table S2**). On the other hand, the free energy barrier for peptide binding (Δ*F_on_*) decreased slightly from 0.69 ± 0.20 kcal/mol in the reweighted profile to 0.61 ± 0.14 kcal/mol in the modified PMF profile (**Table S2**). Furthermore, curvatures of the reweighed (*w*) and modified (*w*\*, no reweighting) free energy profiles were calculated near the peptide bound (“B”) and unbound (“U”) low-energy wells and the energy barrier (“Br”), as well as the ratio of apparent diffusion coefficients calculated from the Pep-GaMD simulations with reweighting (*D*) and without reweighting (modified, *D*\*) (**Table S2**). According to the Kramers’ rate theory, the peptide binding and dissociation were accelerated by 2.55 and 9.47 × 10^3^ times, respectively. Therefore, the reweighted *k_on_* and *k_off_* were calculated as 1.92 ± 1.17 × 10^10^ M^−1^·s^−1^ and 749.19 ± 1.57 s^−1^, respectively.

± 0.32 kcal/mol in the reweighted PMF profile to 1.62 ± 0.32 kcal/mol in the modified PMF profile (**Table S2**). The free energy barrier for peptide binding (Δ*F_on_*) decreased from 1.01 ± 0.18 kcal/mol in the reweighted profile to 0.94 ± 0.22 kcal/mol in the modified PMF profile (**Table S2**). According to the Kramers’ rate theory, the peptide binding actually slowed down by a factor of 0.044 but the dissociation were accelerated by 8.82 × 10^3^ times. Therefore, the reweighted *k_on_* and *k_off_* were calculated as 4.06 ± 2.26×10^10^ M^−1^·s^−1^ and 1.45 ± 1.07×10^3^ s^−1^, respectively. They were comparable to the experimental data^75^ of 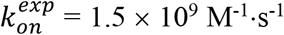 and 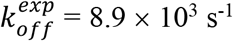 (**Table 2**). Among the three studied peptide systems, the 1CKB system exhibited the highest peptide binding rate constant *k_on_*. This appeared to correlate with the plateau observed in the PMF profile of peptide binding in the 1CKB system (**Fig. 1I**). Although Pep-GaMD could slow down the peptide binding process (1CKA and 1CKB), significant accelerations were achieved in the rate-limiting step of peptide dissociation during Pep-GaMD simulations. The peptides dissociation and binding were more balanced in Pep-GaMD, thereby resulting in overall improved sampling.

### Peptide binding to the SH3 domains was mediated by electrostatic interactions

Next, we examined key residue interactions during peptide binding to the SH3 domains (**Fig. 2**). In the X-ray structures (**Figs. 1A–1C**), a conserved salt-bridge was found between the SH3 domains and bound peptides, i.e., SH3:Asp19 – peptide:Arg10 in the 1SSH structure, SH3:Asp150 - peptide:Lys8 in the 1CKA structure and SH3:Asp150 – peptide:Arg7 in the 1CKB structure. Therefore, the salt-bridge distance between these protein and peptide residues (denoted *d*_D-R/K_) and the peptide backbone RMSD relative to the X-ray structure with the protein aligned were chosen as reaction coordinates to compute 2D PMF profiles. Two low-energy “Bound” and “Intermediate” states were identified from the 2D PMF profiles of all three systems (**Figs. 2A–2C**). In the 1SSH system, (peptide backbone RMSD, *d*_D-R/K_) of the “Bound” and “Intermediate” states were centered around (2.0 Å, 4.2 Å) and (20.0 Å, 5.0 Å), respectively (**Fig. 2A**). In the 1CKA system, (peptide backbone RMSD, d_D-R/K_) of the two states were centered around (2.7 Å, 4.4 Å) and (21.5 Å, 5.2Å) (**Fig. 2B**). In the 1CKB system, (peptide backbone RMSD, *d*_D-R/K_) of the two states were centered around (1.7 Å, 3.7 Å) and (25.0 Å, 8.0 Å) (**Fig. 2C**). Therefore, the salt bridge distances in the “Intermediate” state of the 1SSH and 1CKA structures were similar to those in their “Bound” state, but the salt-bridge distance increased to 8.0 Å in the “Intermediate” state of the 1CKB structure.

**Fig. 2.**
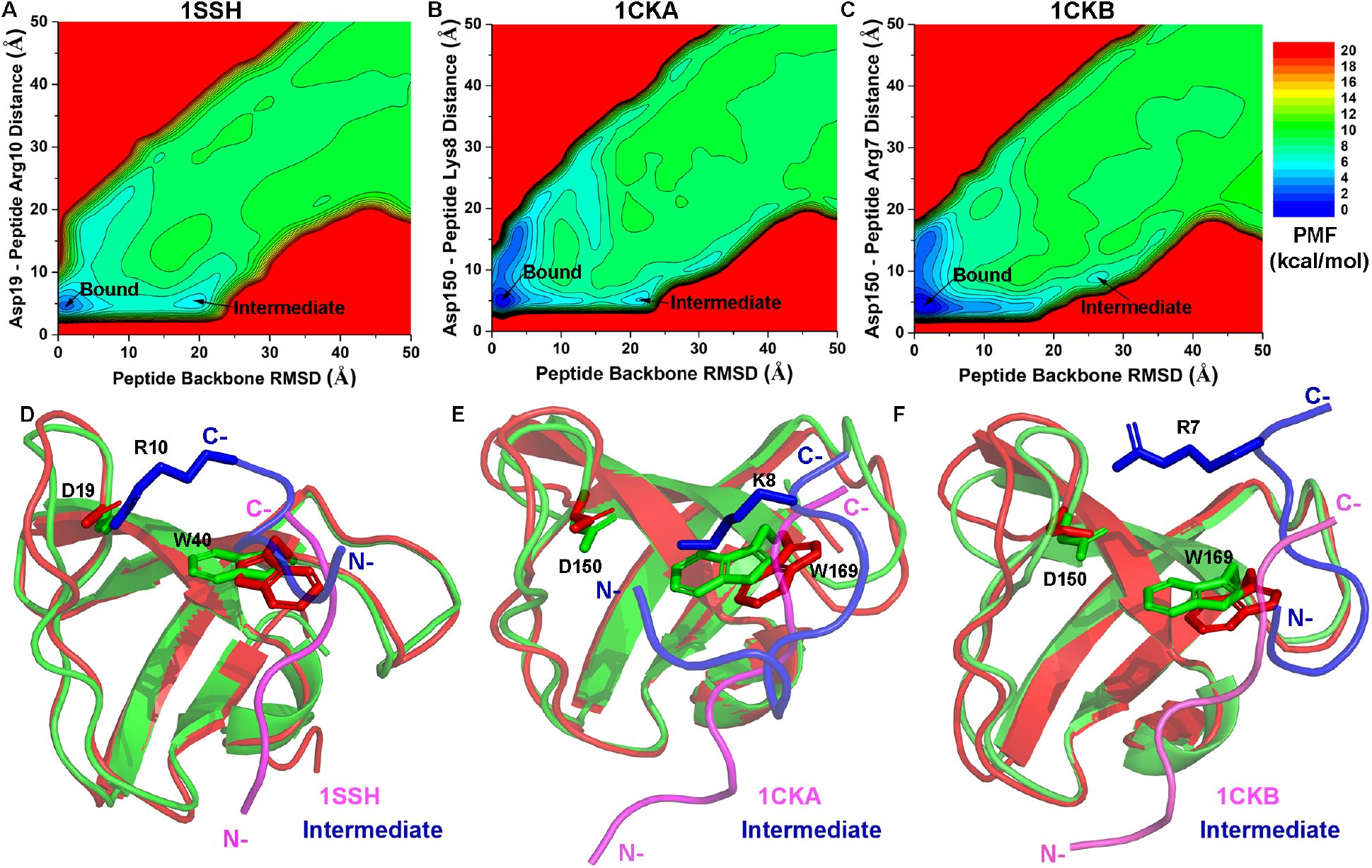
Free energy profiles and low-energy conformational states of peptide binding to the SH3 domains: (A) 2D PMF profiles regarding the peptide backbone RMSD and the distance between protein Asp19 and peptide Arg10 in the 1SSH structure. (B) 2D PMF profiles regarding the peptide backbone RMSD and the distance between protein Asp150 and peptide Lys8 in the 1CKA structure. (C) 2D PMF profiles regarding the peptide backbone RMSD and the distance between protein Asp150 and peptide Arg7 in the 1CKB structure. (D-F) Low-energy intermediate conformations (red) as identified from the 2D PMF profiles of the (D) 1SSH, (E) 1CKA and (F) 1CKB structures, respectively. X-ray structures of the peptide-bound complexes are shown in green and magenta for protein and peptide, respectively. Protein residues Asp19 and Trp40 in the 1SSH structure and Asp150 and Trp169 in the 1CKA and 1CKB structures, and peptide residues Arg10 in the 1SSH structure, Lys8 in the 1CKA structure and Arg7 in the 1CKB structure are highlighted in sticks. The “N” and “C” labels denote the N-terminus and C-terminus of the peptides.

**Figs. 2D–2F** depict the “Intermediate” conformations of the three peptide systems compared with their native “Bound” conformations. While the salt bridge became partially open in the “Intermediate” conformation of the 1CKB system (**Fig. 2F**), it remained fully closed in the “Intermediate” conformations of the 1SSH and 1CKA systems (**Figs. 2D–2E**). In summary, long-range electrostatic interactions played a key role in peptide binding to the SH3 domains. The salt bridge described here appeared to be an anchor that pulled the peptides to their target binding site in the SH3 domains. Similar findings were obtained from previous cMD simulations^19^.

### Peptides bound to the SH3 domains via conformational selection

In order to further explore the mechanism of peptide binding to the SH3 domains, we computed 2D PMF free energy profiles to characterize conformational changes of the three systems upon peptide binding. By closely examining the Pep-GaMD simulation trajectories, we observed the largest conformational changes in protein residue Trp40 in the 1SSH structure or residue Trp169 in the 1CKA and 1CKB structures, which was located at the peptide binding site of the SH3 domains (**Figs. 2D–2F**). Therefore, we calculated 2D reweighted PMF profiles from the Pep-GaMD simulations regarding the peptide backbone RMSD relative to the X-ray structure with the protein aligned and the *d*_D-W_ distance between Asp19-Trp40 in the 1SSH structure or Asp150-Trp169 in the 1CKA and 1CKB structures (**Figs. 3A–3C**). For the 1SSH system, two low-energy states were identified from the 2D PMF profile, including the “Bound” and “Intermediate”, in which the protein residue Trp40 at the peptide-binding site adopted primarily the “Closed” and “Open” conformations, respectively (**Figs. 3A** and **2D**). The peptide backbone RMSD and *d*_D-W_ were centered around (2.0 Å, 6.2 Å) in the “Bound/Closed” state and (9.8 Å, 9.0 Å) in “Intermediate/Open” state (**Fig. 3A**). For the 1CKA system, three low-energy states were identified from the 2D PMF profile, including the “Bound”, “Intermediate” and “Unbound”, in which the protein residue Trp169 adopted primarily the “Closed”, “Open” and “Open” conformations, respectively (**Figs. 3B** and **2E**). The peptide backbone RMSD and *d*_D-W_ were centered around (2.7 Å, 5.9 Å), (13.5 Å, 9.3 Å) and (33.5 Å, 9.3 Å) in the “Bound/Closed”, “Intermediate/Open” and “Unbound/Open” states, respectively (**Fig. 3B**). Three similar low-energy conformational states were identified in the 1CKB system as in the 1CKA system. The peptide backbone RMSD and *d*_D-W_ were centered around (1.7 Å, 5.8 Å), (11.5 Å, 9.3 Å) and (36.0 Å, 8.8 Å) in the “Bound/Closed”, “Intermediate/Open” and “Unbound/Open” states, respectively (**Fig. 3C**).

**Fig. 3.**
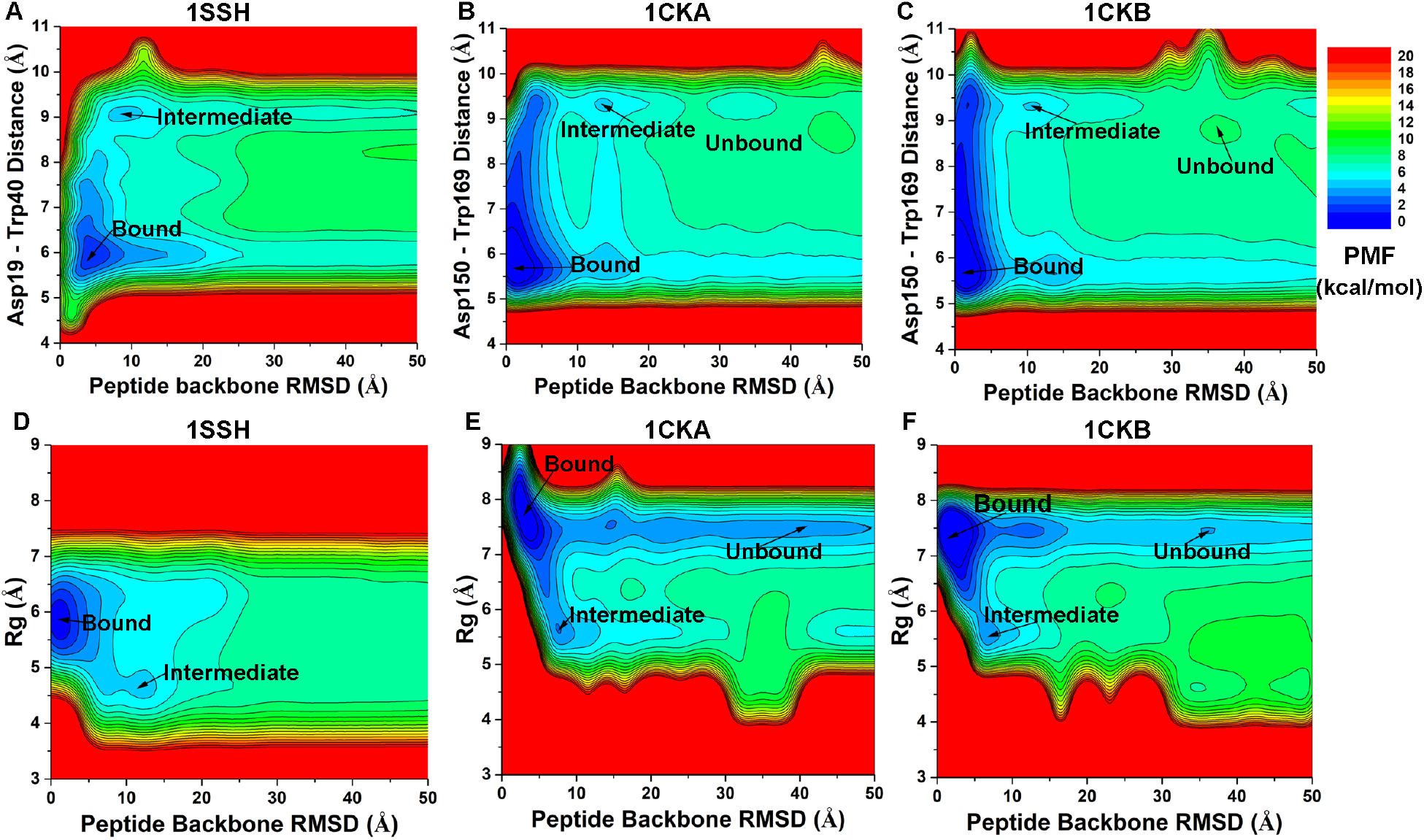
(A) 2D PMF profiles of peptide backbone RMSD and the distance between protein Asp19 and Trp4O calculated from Pep-GaMD simulations of the 1SSH structure. (B-C) 2D PMF profiles of the peptide backbone RMSD and distance between protein Asp150 and Trp169 calculated from Pep-GaMD simulations of the (B) 1CKA and (C) 1CKB structures. (D-F) 2D PMF profiles regarding the peptide backbone RMSD and peptide *R_g_* in the (D) 1SSH, (E) 1CKA and (F) 1CKB structures.

In addition to the target protein, we analyzed conformational changes of the peptides during binding. In this regard, the radius of gyration (*R_g_*) was calculated and monitored for possible conformational changes and even folding and unfolding of the peptides. Then the peptide *R_g_* and backbone RMSDs relative to X-ray structures with SH3 domain aligned were used as reaction coordinates to calculate 2D PMF profiles. Large conformational changes were observed in the three studied peptide systems during the Pep-GaMD simulations, for which large conformational space with a wide range of *R_g_* was sampled for each of the three peptides during binding to SH3 domains (**Figs. 3D–3F**). Therefore, it is important to include the peptide bonded potential energy for applying the first boost potential to account for extremely high flexibility of the peptides. From the reweighted 2D PMF profiles (**Fig. 3D–3F**), we identified a clear low-energy “Bound” state in all three systems, for which the peptide backbone RMSD and *R_g_* in the 1SSH, 1CKA and 1CKB structures were centered around (2.0 Å, 5.9 Å), (2.7 Å, 7.9 Å) and (1.7 Å, 7.5 Å), respectively. Another “Intermediate” low-energy state was also identified in the 1SSH, 1CKA and 1CKB systems, for which the peptide backbone RMSD and *R_g_* were centered around (11.2 Å, 4.7 Å), (8.5 Å, 5.6 Å) and (8.8 Å, 5.6 Å), respectively. Furthermore, a low energy “Unbound” state appeared in the reweighted 2D PMF profiles of the 1CKA (**Fig. 3E**) and 1CKB (**Fig. 3F**), but not in the reweighted 2D PMF profile of the 1SSH system (**Fig. 3A**). Nevertheless, the “Unbound” state was found in the 2D PMF profiles without energetic reweighting for all three systems, similarly for the “Bound” and “Intermediate” states (**Fig. S2**). This suggested that complete peptide binding processes were successfully sampled in the Pep-GaMD simulations, despite differences in the free energy profiles. More importantly, the peptides sampled a large conformational space with a wide range of *R*g in the “Unbound” state, but only a subset of the conformations were chosen upon binding to the target protein. Therefore, peptide binding to the SH3 domains followed predominantly a “conformational selection” mechanism.

## Conclusions

A new computational method called Pep-GaMD has been developed for efficient enhanced sampling and free energy and kinetics calculations of peptide binding. Pep-GaMD works by selectively boosting the essential potential energy of the peptide to effectively model its high flexibility. Microsecond Pep-GaMD simulations have allowed us to capture repetitive peptide dissociation and binding as demonstrated on binding of three model peptides to the SH3 domains. Pep-GaMD has thus enabled highly efficient free energy and kinetics calculations of peptide binding.

Pep-GaMD appears to be more efficient and/or easier to use for enhanced sampling of peptide binding as compared with existing methods, including the cMD^26^, replica exchange^33, 34, 37^, weighted ensemble^23^, Markov state models^36^ and metadynamics^37^. In particular, 200 μs Anton cMD simulations at elevated temperature (400 K) were performed to capture repetitive dissociation and binding of an IDP peptide^26^. A number of replica simulations were needed to model peptide binding using the replica exchange algorithm^33, 34, 37^, while an independent microsecond Pep-GaMD simulation was able to capture multiple events of peptide binding and unbinding. A total of ~120 μs cMD simulations were needed to calculate peptide binding rate constants using the weighted ensemble approach^23^. Multi-ensemble Markov models that were used to characterize kinetics of the PMI peptide binding to the MDM2 protein were constructed using ~500 μs cMD and ~50 μs Hamiltonian replica exchange simulations combined^76^. Bias-exchange metadynamics simulations and a total of 27 μs infrequent metadynamics simulations were applied to calculate the p53-MDM2 binding free energy and kinetic rates, respectively^37^, but they required predefined, carefully chosen CVs with expert knowledge of the studied system. Moreover, the predefined CVs could potentially lead to constraints on the peptide binding pathway and conformational space to be sampled. When important CVs were missed, the simulations could suffer from the “hidden energy barrier” problem and slow convergence^78^. Overall, the above methods were still computationally expensive, requiring tens to hundreds of microsecond simulations to characterize peptide binding thermodynamics and kinetics. In this context, Pep-

GaMD that has enabled such calculations through only microseconds simulations provides a highly efficient and easy-to-use approach to enhanced sampling of peptide binding, although testing on more peptide binding systems are still needed. Pep-GaMD has already been implemented in the GPU version of AMBER 20 simulation package, which should facilitate further testing and usage of the method^58^ (see **Methods** in **Supplementary Material**).

Pep-GaMD simulations showed that long-range electrostatic effects played an important role in peptide binding to the SH3 domains. The electrostatic interactions were also identified in earlier cMD^19^ and NMR studies^76, 79^. For example, multiple short cMD simulations (aggregated to 0.8 μs) were performed to capture binding of a PRM peptide to the C-CRK N-terminal SH3 domain^19^. The simulations suggested that long-range electrostatic effects played a major role in the peptide diffusion and facilitated formation of a transient peptide-protein complex. Skrynnikov et al.^76, 79^ combined cMD and NMR experiments to investigate electrostatic interactions involved in the formation of the electrostatic encounter complex. With improved force field parameters and long cMD simulations up to 3 μs, binding of the Sos peptide to the C-CRK N-terminal SH3 domain was successfully captured. However, peptide dissociation was still not observed in these long cMD simulations. In contrast, microsecond Pep-GaMD simulations here captured both binding and unbinding of peptides, further supporting the important role of electrostatic interactions in forming the “intermediate” and “bound” states of the peptide-protein systems (**Fig. 2**).

Pep-GaMD shall be of wide applicability for studies of many other peptide-protein interactions^1^ other than peptide binding to the SH3 domains. Beyond accurate peptide binding free energy (thermodynamics) calculations, we have also successfully derived the dissociation and binding kinetic rate constants of peptide binding to the SH3 domains. While peptide dissociation has been accelerated by ~4-5 orders of magnitude, peptide binding in two of the three studies systems have been surprisingly slowed down in this study (Table 2). To facilitate peptide rebinding in future studies, one can include multiple peptides in the system within their solubility limits, an approach that has been shown to accelerate ligand binding in the recently developed LiGaMD method^49^. These developments are expected to further improve the Pep-GaMD method for applications in enhanced sampling of peptide-protein interactions.

## Acknowledgements

We appreciate the help of Prof. David Case for accessing the AMBER git repository to develop our new simulation algorithms. This work used supercomputing resources with allocation award TG-MCB180049 through the Extreme Science and Engineering Discovery Environment (XSEDE), which is supported by National Science Foundation grant number ACI-1548562, and project M2874 through the National Energy Research Scientific Computing Center (NERSC), which is a U.S. Department of Energy Office of Science User Facility operated under Contract No. DE-AC02-05CH11231, and the Research Computing Cluster at the University of Kansas. This work was supported in part by the National Institutes of Health (R01GM132572) and the startup funding in the College of Liberal Arts and Sciences at the University of Kansas.

## Supplementary Material

Detailed description of the GaMD and energetic reweighting methods, Tables S1-S2 and **Figs. S1-S2** are provided in the **Supplementary Material**.

## Data Availability Statement

The data that supports the findings of this study are available within the article and its supplementary material. The all-atom simulation files are available from the corresponding author upon reasonable request.

