## Supplementary Material for "Peptide Gaussian accelerated molecular dynamics (Pep-GaMD): Enhanced sampling and free energy and kinetics calculations of peptide binding"

### Methods

#### Gaussian accelerated molecular dynamics (GaMD)

Consider a system with $N$ atoms at positions $r \equiv \{\vec{r}_1, \cdots, \vec{r}_N\}$. When the system potential $V(r)$ is lower than a reference energy $E$, the modified potential $V^*(r)$ of the system is calculated as:

$$V^*(r) = V(r) + \Delta V(r),$$

$$\Delta V(r) = \begin{cases} \frac{1}{2}k\big(E - V(r)\big)^2, & V(r) < E \\ 0, & V(r) \geq E \end{cases} \quad \text{(S1)}$$

where $k$ is the harmonic force constant. The two adjustable parameters $E$ and $k$ are automatically determined based on three enhanced sampling principles[1]. The reference energy needs to be set in the following range:

$$V_{max} \leq E \leq V_{min} + \frac{1}{k}, \quad \text{(S2)}$$

where $V_{max}$ and $V_{min}$ are the system minimum and maximum potential energies. To ensure that Eqn. (S2) is valid, $k$ has to satisfy: $k \leq \frac{1}{V_{max}-V_{min}}$ Let us define $\equiv k_0 \cdot \frac{1}{V_{max}-V_{min}}$, then $0 < k_0 \leq 1$. The standard deviation of $\Delta V$ needs to be small enough (i.e., narrow distribution) to ensure proper energetic reweighting[2]: $\sigma_{\Delta V} = k\big(E - V_{avg}\big)\sigma_V \leq \sigma_0$ where $V_{avg}$ and $\sigma_V$ are the average and standard deviation of the system potential energies, $\sigma_{\Delta V}$ is the standard deviation of $\Delta V$ with $\sigma_0$ as a user-specified upper limit (e.g., $10k_B$T) for proper reweighting. When $E$ is set to the lower bound $E$=$V_{max}$, $k_0$ can be calculated as:

$$k_0 = \min(1.0, k_0') = \min\,(1.0, \frac{\sigma_0}{\sigma_V} \cdot \frac{V_{max}-V_{min}}{V_{max}-V_{avg}}). \quad \text{(S3)}$$

Alternatively, when the threshold energy $E$ is set to its upper bound $E = V_{min} + \frac{1}{k}$, $k_0$ is set to:

$$k_0 = k_0'' \equiv (1 - \frac{\sigma_0}{\sigma_V})\frac{V_{max}-V_{min}}{V_{avg}-V_{min}}, \quad \text{(S4)}$$

if $k_0''$ is found to be between *0* and *1*. Otherwise, $k_0$ is calculated using Eqn. (S3).

#### Energetic reweighting of GaMD simulations

For energetic reweighting of GaMD simulations to calculate potential of mean force (PMF), the probability distribution along a reaction coordinate is written as $p^*(A)$. Given the boost potential $\Delta V(r)$ of each frame, $p^*(A)$ can be reweighted to recover the canonical ensemble distribution, $p(A)$, as:

$$p\big(A_j\big) = p^*\big(A_j\big)\frac{\langle e^{\beta\Delta V(r)}\rangle_j}{\sum_{i=1}^{M}\langle p^*(A_i)e^{\beta\Delta V(r)}\rangle_i}, \; j = 1, \ldots, M, \quad \text{(S5)}$$

where $M$ is the number of bins, $\beta = k_BT$ and $\langle e^{\beta\Delta V(r)}\rangle_j$ is the ensemble-averaged Boltzmann factor of $\Delta V(r)$ for simulation frames found in the $j^{th}$ bin. The ensemble-averaged reweighting factor can be approximated using cumulant expansion:

$$\langle e^{\beta\Delta V(r)}\rangle = exp\left\{\sum_{k=1}^{\infty}\frac{\beta^k}{k!}C_k\right\}, \quad \text{(S6)}$$

where the first two cumulants are given by:

$$C_1 = \langle \Delta V \rangle, \\ C_2 = \langle \Delta V^2 \rangle - \langle \Delta V \rangle^2 = \sigma_v^2. \tag{S7}$$

The boost potential obtained from GaMD simulations usually follows near-Gaussian distribution[3]. Cumulant expansion to the second order thus provides a good approximation for computing the reweighting factor[1, 2]. The reweighted free energy $F(A) = -k_B T \ln p(A)$ is calculated as:

$$F(A) = F^*(A) - \sum_{k=1}^{2} \frac{\beta^k}{k!} C_k + F_c, \tag{S8}$$

where $F^*(A) = -k_B T \ln p^*(A)$ is the modified free energy obtained from GaMD simulation and $F_c$ is a constant.

**Implementation of peptide Gaussian accelerated molecular dynamics (Pep-GaMD)**

Peptide Gaussian accelerated molecular dynamics (Pep-GaMD) is currently implemented in the GPU version of AMBER 20[4], but should be transferable to other molecular dynamics programs as well. Pep-GaMD provides enhanced sampling of peptide binding and unbinding to proteins. Following is a list of the input parameters for a Pep-GaMD simulation:

***igamd*** Flag to apply boost potential
= **0** (default) no boost is applied
= **1** boost on the total potential energy only (GaMD_Tot)
= **2** boost on the dihedral energy only (GaMD_Dih)
= **3** dual boost on both dihedral and total potential energy (GaMD_Dual)
= **4** boost on the non-bonded potential energy only (GaMD_NB)
= **5** dual boost on both dihedral and non-bonded potential energy (GaMD_NB_Dual)
= **10** boost on non-bonded potential energy of selected region (defined by timask1 and scmask1) as for a ligand (LiGaMD)
= **11** dual boost on both non-bonded potential energy of the bound ligand and the remaining potential energy of the entire system (LiGaMD_Dual)
= **14** boost on the total potential energy of selected region (defined by timask1 and scmask1) as for a peptide (Pep-GaMD)
= **15** dual boost on both the peptide essential interaction potential energy and the remaining potential energy of the entire system (Pep-GaMD_Dual)

***iE*** Flag to set the threshold energy $E$ for applying all boost potentials
= **1** (default) set the threshold energy to the lower bound $E = V_{max}$
= **2** set the threshold energy to the upper bound $E = V_{min} + (V_{max} - V_{min})/k_0$

***iEP*** Flag to overwrite *iE* and set the threshold energy $E$ for applying the first boost potential in dual-boost schemes
= **1** (default) set the threshold energy to the lower bound $E = V_{max}$
= **2** set the threshold energy to the upper bound $E = V_{min} + (V_{max} - V_{min})/k_0$

***iED*** Flag to overwrite *iE* and set the threshold energy $E$ for applying the second boost potential in dual-boost schemes
= **1** (default) set the threshold energy to the lower bound $E = V_{max}$
= **2** set the threshold energy to the upper bound $E = V_{min} + (V_{max} - V_{min})/k_0$

***ntcmdprep*** The number of preparation conventional molecular dynamics steps. This is used for system equilibration and the potential energies are not collected for calculating their statistics. The default is 200,000 for a simulation with 2 *fs* timestep.

***ntcmd*** The number of initial conventional molecular dynamics simulation steps used to calculate the maximum, minimum, average and standard deviation of the system potential energies (i.e., $V_{max}$, $V_{min}$, $V_{avg}$, $\sigma_V$). The default is 1,000,000 for a simulation with 2 *fs* timestep.

***ntebprep*** The number of preparation biasing molecular dynamics simulation steps. This is used for system equilibration after adding the boost potential and the potential statistics (i.e., $V_{max}$, $V_{min}$, $V_{avg}$, $\sigma_V$) are not updated during these steps. The default is 200,000 for a simulation with 2 fs timestep.

***nteb*** The number of biasing molecular dynamics simulation steps. Potential statistics ($V_{max}$, $V_{min}$, $V_{avg}$, $\sigma_V$) are updated between the ***ntebprep*** and ***nteb*** steps and used to calculate the GaMD acceleration parameters, particularly $E$ and $k_0$. The default is 1,000,000 for a simulation with 2 *fs* timestep. A greater value may be needed to ensure that the potential statistics and GaMD acceleration parameters level off before running production simulation between the ***nteb*** and ***nstlim*** (total simulation length) steps. Moreover, ***nteb*** can be set to ***nstlim***, by which the potential statistics and GaMD acceleration parameters are updated adaptively throughout the simulation. This in some cases provides more appropriate acceleration.

***ntave*** The number of simulation steps used to calculate the average and standard deviation of potential energies. This variable has already been used in AMBER. The default is set to 50,000 for GaMD simulations. It is recommended to be updated as about 4 times of the total number of atoms in the system. Note that ***ntcmd*** and ***nteb*** need to be multiples of ***ntave***.

***irest_gamd*** Flag to restart GaMD simulation
**= 0** (default) new simulation. A file "gamd-restart.dat" that stores the maximum, minimum, average and standard deviation of the potential energies needed to calculate the boost potentials (depending on the ***igamd*** flag) will be saved automatically after GaMD equilibration stage.
**= 1** restart simulation (***ntcmd*** and ***nteb*** are set to 0 in this case). The "gamd-restart.dat" file will be read for restart.

***sigma0P*** The upper limit of the standard deviation of the first potential boost that allows for accurate reweighting. The default is 6.0 (unit: kcal/mol).

***sigma0D*** The upper limit of the standard deviation of the second potential boost that allows for accurate reweighting in dual-boost simulations (e.g., ***igamd*** = 2, 3, 5 and 11). The default is 6.0 (unit: kcal/mol).

***timask1*** Specifies atoms of the bound ligand in ambmask format. This variable has already been used in AMBER. The default is an empty string.

***scmask1*** Specifies atoms of the bound ligand that will be described using soft core in ambmask format. This variable has already been used in AMBER. The default is an empty string.

Example input parameters used in Pep-GaMD_Dual simulations include the following:

```
icfe = 1, ifsc = 1, gti_cpu_output = 0,gti_add_sc = 1,
```

```
timask1 = ':1-3', scmask1 = ':1-3',
timask2 = '', scmask2 = '',

igamd = 15, iEP = 2, iED = 1, irest_gamd = 0,
ntcmd = 1000000, nteb = 1000000, ntave = 50000,
ntcmdprep = 200000, ntebprep = 200000,
sigma0P = 6.0, sigma0D = 6.0,
```

The Pep-GaMD algorithm is summarized as the following:

```
Pep-GaMD {
  If (irest_gamd == 0) then
   For i = 1, …, ntcmd // run initial conventional molecular dynamics
     If (i >= ntcmdprep) Update Vmax, Vmin
     If (i >= ntcmdprep && i%ntave ==0) Update Vavg, sigmaV
   End
   Save Vmax,Vmin,Vavg,sigmaV to "gamd_restart.dat" file
   Calc_E_k0(iE,sigma0,Vmax,Vmin,Vavg,sigmaV)

   For i = ntcmd+1, …, ntcmd+nteb // Run biasing molecular dynamics simulation steps
      deltaV = 0.5*k0*(E-V)**2/(Vmax-Vmin)
      V = V + deltaV
      If (i >= ntcmd+ntebprep) Update Vmax, Vmin
      If (i >= ntcmd+ntebprep && i%ntave ==0) Update Vavg, sigmaV
      Calc_E_k0(iE,sigma0,Vmax,Vmin,Vavg,sigmaV)
   End
   Save Vmax,Vmin,Vavg,sigmaV to "gamd_restart.dat" file
  else if (irest_gamd == 1) then
   Read Vmax,Vmin,Vavg, sigmaV from "gamd_restart.dat" file
  End if


  For i = ntcmd+nteb+1, …, nstlim // run production simulation
   deltaV = 0.5*k0*(E-V)**2/(Vmax-Vmin)
   V = V + deltaV
  End
}
```

```
Subroutine Calc_E_k0(iE,sigma0,Vmax,Vmin,Vavg,sigmaV) {
if iE = 1 :
        E = Vmax
        k0' = (sigma0/sigmaV) * (Vmax-Vmin)/(Vmax-Vavg)
        k0 = min(1.0, k0')
else if iE = 2 :
        k0" = (1-sigma0/sigmaV) * (Vmax-Vmin)/(Vavg-Vmin)
        if 0 < k0" <= 1 :
                k0 = k0"
                E = Vmin + (Vmax-Vmin)/k0
        else
                E = Vmax
                k0' = (sigma0/sigmaV) * (Vmax-Vmin)/(Vmax-Vavg)
                k0 = min(1.0, k0')
        end
end
}
```

**Table S1.** The peptide binding and unbinding time periods ($\tau_B$ and $\tau_U$) recorded from Pep-GaMD simulations of the peptide-SH3 domain binding systems.

| system | | $\tau_B$ (ns) | $\tau_U$ (ns) |
|---|---|---|---|
| 1ssh | sim1 | 195.39,9.35,229.59,59.09 | 28.73,65.43,144.33,204.51 |
| | sim2 | 256.95,257.18,34.20 | 20.98,273.05 |
| | sim3 | 68.17,164.93 | 48.56,415.87 |
| 1cka | sim1 | 87.77,5.35,10.13,45.83,38.17 | 24.86,237.34,156.54,287.68 |
| | sim2 | 16.87,98.50,22.34 | 351.57,73.41 |
| | sim3 | 102.14,38.08 | 187.64,654.58 |
| 1ckb | sim1 | 15.73,51.08,94.39,36.71 | 39.44,541.70,40.70 |
| | sim2 | 45.64,100.87,216.21 | 379.04,264.54 |
| | sim3 | 23.62,34.80,141.4 | 108.52,80.76 |

**Table S2.** Energy barriers of peptide-SH3 dissociation ("off") and binding ("on") calculated from the reweighed (***ΔF***) and modified (no reweighting, ***ΔF****) free energy profiles, curvatures of the reweighed ($w$) and modified ($w*$) free energy profiles near the guest Bound ("B"), Barrier ("Br") and Unbound ("U") states, and the ratio of apparent diffusion coefficients calculated from the Pep-GaMD simulations without reweighting (modified, $D*$) and with reweighting ($D$).

| system | Δ***F*** (kcal/mol) | | Δ***F**** (kcal/mol) | | ***w*** | | | ***w**** | | | ***D*/D*** | |
|---|---|---|---|---|---|---|---|---|---|---|---|---|
| | **Off** | **On** | **Off** | **On** | **B** | **Br** | **U** | **B** | **Br** | **U** | **Off** | **On** |
| 1SSH | 8.16±0.16 | 0.69±0.20 | 2.40±0.32 | 0.61±0.14 | 0.65±0.074 | 0.26±0.10 | 0.041±0.023 | 0.47±0.028 | 0.12±0.038 | 0.0052±0.0013 | 0.52±0.030 | 1.22±0.54 |
| 1cka | 7.42±0.64 | 2.02±1.12 | 1.35±0.084 | 1.17±0.04 | 0.74±0.024 | 0.54±0.25 | 0.12±0.13 | 0.72±0.041 | 0.22±0.054 | 0.034±0.034 | 2.48±1.99 | 2.17±2.23 |
| 1ckb | 7.84±0.32 | 1.01±0.18 | 1.62±0.32 | 0.94±0.22 | 0.96±0.22 | 0.31±0.075 | 0.030±0.018 | 0.75±0.19 | 0.12±0.084 | 0.010±0.003 | 0.81±0.019 | 0.54±0.38 |

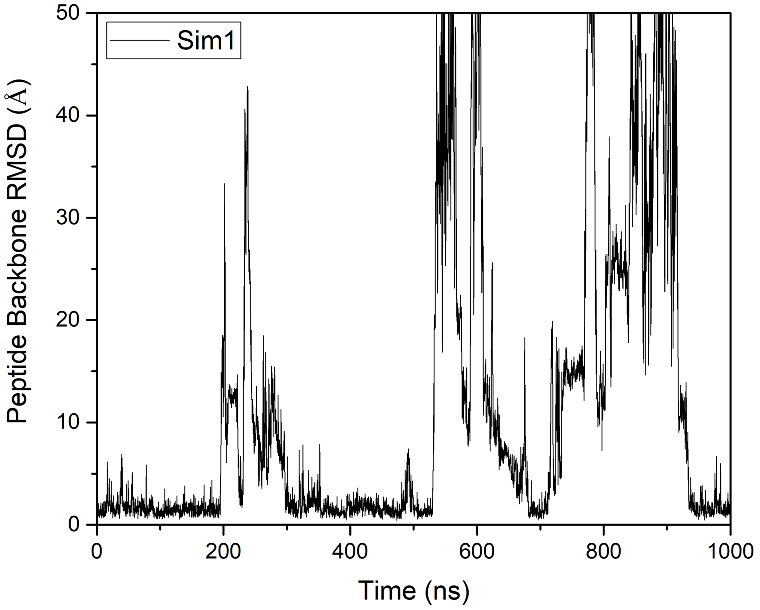


**Fig S1.** Time course of peptide backbone RMSDs relative to X-ray structures with the protein aligned calculated from Sim1 Pep-GaMD simulations of the 1SSH.

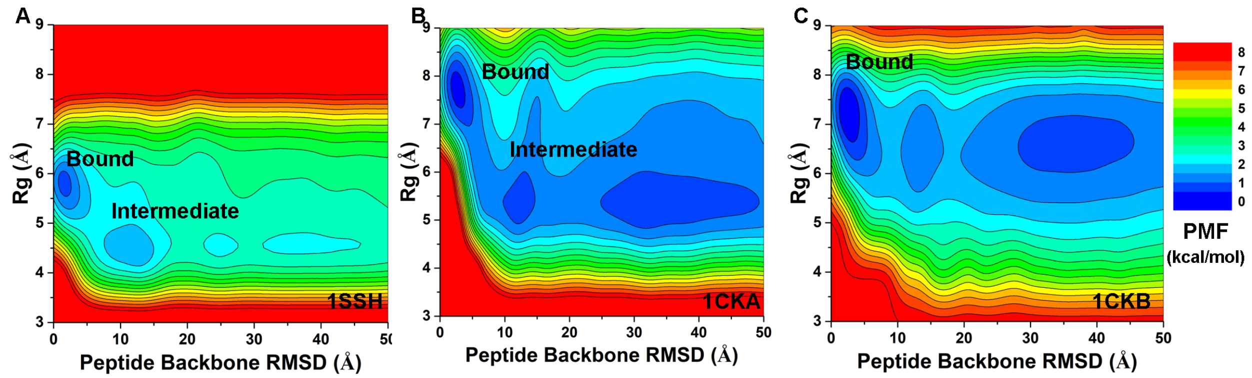


**Fig. S2.** Un-reweighted 2D PMF free energy profiles regarding the peptide backbone RMSD and peptide $R_g$ calculated from Pep-GaMD simulations of the (A) 1SSH, (B) 1CKA and (C) 1CKB structures.
